# A nucleic acid host factor enables optimal phage replication in *Escherichia coli*

**DOI:** 10.64898/2026.06.19.733364

**Authors:** Alice CZ Collins, Marcel Sprenger, Josh McQuail, Sebastian Krautwurst, Kai Papenfort, Sivaramesh Wigneshweraraj

## Abstract

Host acquisition by bacteriophages (phages) often entails modulation, appropriation, or inhibition of components and processes central to bacterial gene expression. Small non-coding RNAs (sRNAs) are major regulators of RNA fate and frequently rely on the conserved RNA chaperone Hfq to engage their cognate targets. Although phages are known to encode specialised proteins and sRNAs to manipulate host gene expression, it has remained unclear whether they also co-opt host-encoded sRNAs for their own regulatory needs. We show that transcriptome-wide Hfq-mediated RNA–RNA interactions are broadly destabilised during T2 phage infection of *Escherichia coli*. We further demonstrate that the conserved bacterial sRNA ArcZ is co-opted by T2 to promote expression of a conserved phage operon that includes a protein which inhibits a bacterial restriction–modification system. ArcZ achieves this by preventing RNase E-mediated degradation of the transcript originating from the phage operon. Our study provides the first evidence of an evolutionary strategy in which a phage leverages a nucleic acid host factor to fulfil its own gene expression requirements.

## Introduction

The regulation of bacterial gene expression occurs primarily at two key stages. First, transcriptional control is mediated by regulatory proteins that influence the activity of RNA polymerase (RNAP), the enzyme responsible for synthesizing RNA. These proteins can act as activators or repressors, fine-tuning the initiation and rate of transcription in response to cellular cues. Second, post-transcriptional regulation involves small non-coding RNA molecules (sRNA), which modulate the stability, translation efficiency, and degradation of their cognate target RNA molecules. Some sRNA interact with other sRNAs, a process referred to as ‘sponging’ and represents a regulatory mechanism for sequestering sRNA or promoting their degradation (1). In many bacteria, a key mediator of post-transcriptional regulation is the RNA-binding protein Hfq, which acts as an RNA chaperone to facilitate interactions between sRNAs and their cognate RNA targets (1,2).

Viruses that infect bacteria, bacteriophages (phages), have evolved innovative ways to co-opt host proteins to benefit their replication and progeny development (3). Unsurprisingly, host processes associated with RNA metabolism are also targeted. Phages also produce small proteins which either inhibit host RNAP from utilising host promoters or direct it to exclusively utilise phage promoters (4). Other phage-encoded proteins modulate the activity of host RNases, compromising global RNA stability whilst benefiting phage RNA maturation and stability (5–8). Hfq was originally identified as the host protein required for RNA phage Qβ replication, where Hfq is required to induce structural changes in phage RNA, allowing engagement with the replication machinery (9). Many other phages also rely on Hfq for their post-transcriptional regulation. Several phages have been shown to encode Hfq-dependent sRNAs that regulate host transcripts or ‘sponge’ host sRNAs, both of which serve to benefit the phage (reviewed in (10) and recent examples (11,12)). In contrast, the role of host sRNAs in the post-transcriptional regulation of phage transcripts remains poorly understood. To address this gap in our understanding of phage-bacteria interactions, we mapped the genome-wide Hfq-mediated RNA-RNA interactions during replication of the prototypical *Escherichia coli* phage T2. We uncovered a role for a conserved bacterial sRNA, ArcZ, in the positive post- transcriptional regulation of a phage-derived transcript, which encodes proteins involved in counter-phage defence mechanism. Thus, this study conceptually extends our understanding of phage-bacteria interaction in which a host-derived nucleic acid factor aids phage replication.

## Materials and methods

### Bacterial strains and plasmids

The bacterial strains and plasmids used in this study are summarised in **Supplementary Data 1**. Bacteria were grown in lysogeny broth LB at 37 °C. Colony forming units (CFU) were measured to enumerate the proportion of viable cells by serial dilution of the culture and plating on LB agar plates. Bacterial cells were made chemically competent for transformation of plasmids following standard laboratory procedures. Antibiotics were used at the following concentrations: carbenicillin 100 μg/ml, chloramphenicol 35 μg/ml, zeocin 100 μg/ml, kanamycin 50 μg/ml. λ Red recombination was used to delete *arcZ* from the Δ*mcrC* strain as previously described (13). All plasmids used in **Figure 3** were constructed by Genewiz PriorityGENE service using pBAD18 as the backbone vector. All nucleotide sequences corresponding to the construct indicated in **Figure 3** were placed under the control of the pBAD promoter. pArcZ was constructed using pKF68-3 (14) as the backbone, with *Salmonella sdsR* changed for *E. coli arcZ* using PCR products amplified from gDNA and constructed by Gibson assembly (15).

### T2 phage infection assays

Overnight bacterial cultures for the RIL/RNA-seq experiments were grown for 2 hrs to OD_600_ ∼0.3 at 37 °C in LB. T2 was added at multiplicity of infection (MOI) ∼20 and samples for analyses were collected before phage addition and at the indicated time points (see Results). For the assay in **Supplementary Figure 1C**, bacteria were grown in flasks to OD_600_ ∼0.2 and used to inoculate LB -/+ T2 (at MOI ∼20) in flat-bottomed 96-well plate. The plate was put in a SPECTROstar Nano microplate reader (BMG LABTECH), and OD_600_ readings were taken every 15 min.

### Plaque forming units (PFU) measurements

T2 lysate (100 μl) was mixed with 100 μl of recipient bacterial culture (OD_600_ ∼0.34) as indicated in the figures and incubated at room temperature for 5 mins. Phage top agar (PTA; 20 g LB broth, 4 g agar, 10 mM CaCl_2_) was added; if the recipient bacterial cells contained plasmids for expression of Arn proteins, ArcZ or Hfq (see Results), 0.2% final concentration of L-arabinose (for pBAD-based plasmids) or 0.1 mM IPTG (for pArcZ) was added to PTA and poured onto plates made with phage bottom agar (PBA: 20 g LB broth, 7 g agar, 10 mM CaCl_2_). The plates were incubated overnight at 37 °C. PTA was collected and added to phage buffer (PHB; 50 mM Tris-HCl pH8.0, 1 mM NaCl, 1 mM MgSO_4_, 4 mM CaCl_2_) vortexed before centrifuging for 10 min at ∼4000 RPM. The supernatant (cleared lysate) was then filtered through 0.22 μm filter. The filtered lysate was used to enumerate PFU as follows: A bacterial lawn was prepared by mixing WT bacteria (100 μl, OD_600_ ∼0.34) with PTA and pouring the mixture onto PBA plates. The filtered phage lysates were serially diluted and spotted onto the solidified PTA of the prepared plates and incubated overnight at 37 °C. PFUs were counted and quantified as PFU/ml.

### RIL-seq

RIL-seq experiments and data analysis were performed as previously described (16). Briefly, *E. coli hfq*::3X3XFLAG strains were grown in LB at 37 °C to OD_600_ ∼0.2 and either infected with T2 phage MOI 20 or left uninfected. Three biological replicates was used for each condition. Samples equivalent to 50 OD_600_ units were collected just prior to T2 infection, and additionally 20 and 60 mins after T2 infection induction. The samples were UV-crosslinked followed by cell lysis and co-immunoprecipitation using a monoclonal ɑ-3XFLAG antibody (Sigma, F1804). Recovered RNA was trimmed by RNase A/T1 treatment, and proximal RNAs were ligated using T4 RNA ligase. After protein digestion using proteinase K, RNA was extracted, fragmented and TurboDNase digested. Total RNA concentrations were then determined via Nano Chip. Ribosomal RNA was depleted using rRNA-specific biotinylated probes (17). Co-immunoprecipitated RNA was mixed with 1X SSC, 1 mM EDTA and an oligonucleotide mix (5.8 nM for 16S and 23S oligos; 11.6 nM for 5S oligos). The mixture was denatured, cooled down and incubated with streptavidin beads (ThermoFisher; 65001) at room temperature and 50 °C, each step 5 min, respectively. rRNA depleted RNA was purified using the Agencourt AMPure XP kit (Beckman Coulter Genomics) and subjected to cDNA library preparation. The NEBNext Ultra II Q5 polymerase was used for library amplification. The DNA libraries were sequenced on a NextSeq1000 system with 200-nt read length in paired-end mode and chimeric reads were aligned to the *E. coli* strain MG1655 and T2 phage reference genomes (NCBI accession numbers NC_000913.3 and NC_054931, respectively). For further downstream analysis and interpretation, only interactions that were detected with at least 10 chimeras in any one of the three biological replicates were included (FDR-adjusted *p*-value ≤ 0.05) (18). Chimera data can be found in **Supplementary Data 2** and **Supplementary Data 3**.

### RNA-seq

Samples for RNA-seq were collected at the same time as for RIL-seq. Transcription was stopped by adding 0.2 volumes of stop mix (95% ethanol, 5% [vol/vol] phenol). Total RNA was isolated using the TRIzol™ LS reagent (Sigma), treated with Turbo DNase (Thermo Fisher Scientific) and the RNA integrity was confirmed using a Bioanalyzer (Agilent). Ribosomal RNA was depleted using rRNA-specific biotinylated probes as described above. rRNA depleted RNA was purified using the Agencourt AMPure XP kit (Beckman Coulter Genomics) and fragmented for 5 min at 75°C using the NEBNext® Magnesium RNA Fragmentation Module (NEB). cDNA libraries were prepared using the NEBNext Small RNA Library Prep Set for Illumina (NEB; E7300L) according to the manufacturer’s instructions. cDNA library quality was confirmed on an Agilent 2100 Bioanalyzer, and pooled cDNA libraries were sequenced on a NextSeq1000 system with 100-nt read length in paired-read mode. Demultiplexed raw reads were trimmed for quality and 3’ adaptors and mapped to the *E. coli* strain MG1655 and T2 phage reference genomes (NCBI accession numbers NC_000913.3, NC_054931) using “RNA-Seq Analysis” tool of CLC Genomics Workbench (Qiagen) with standard parameter settings. Genes with a fold-change ≥ ±2 and an FDR-adjusted *p*-value ≤ 0.05 were defined as differentially expressed (**Supplementary Data 4**).

### Proteomics

For total proteome analysis of T2 infected WT and Δ*arcZ* bacteria, 20 ml of sample was collected 60 mins after T2 infection, washed twice in cold phosphate buffered saline, pelleted by centrifugation and flash frozen in liquid nitrogen. The pellets were processed and the data analysed as previously described (13). Analysed data used both a Uniprot *E. coli* FASTA database (UP000000625, *E. coli* strain K12, ID83333, 4402 entries, last modified October 27^th^, 2022, downloaded January 11th, 2023) and a Uniprot Phage T2 FASTA database (UP000249673, 278 entries, downloaded April 24th, 2025) filtering out contaminants and reverse sequences. Only proteins quantified with at least 2 razor peptides were considered for further analysis. The following modifications were included into the search parameters: Carbamidomethylation (C, 57.0215), TMTpro (K, 304.2072) as fixed modifications; Oxidation (M, 15.9949), Acetylation (protein N-terminus, 42.0106), TMTpro (peptide N-terminus, 304.2072) as variable modifications. For the full scan (MS1) a mass error tolerance of 20 PPM and for MS/MS (MS2) spectra of 20 PPM was set. For protein digestion, ’trypsin’ was used as protease with an allowance of maximum 2 missed cleavages requiring a minimum peptide length of 7 amino acids. The false discovery rate on peptide and protein level was set to 0.01. Limma analysis for Δ*arcZ* as compared to WT infection with T2 phage can be found in **Supplementary Data 5**.

### Immunoblotting

Whole cell extracts for immunoblotting were prepared as follows: Overnight cultures containing the reporter plasmids/strain combination indicated in **Figure 3** (also see **Supplementary Table S1**) and corresponding ‘empty’ control plasmid were used to inoculate 25 ml of LB and grown at 37 °C to OD600 ∼0.3. L-arabinose was added to the cultures at a final concentration of 0.2% (v/v) to induce expression of the constructs under the pBAD promoter and sampled at the time points indicated in **Figure 3**. A sample just prior to L-arabinose addition served as the T0 time point. For the T0, 3 ml of culture was collected, whilst subsequent time points, 2 ml of sample. For **Supplementary Figure 2B**, overnight cultures of MG1655 with *hfq* 3XFLAG-tagged (also see **Supplementary Data 1**) were used to inoculate 25 ml of LB and grown at 37 °C to OD600 ∼0.3. T2 phage was added to infect one flask at MOI 20, leaving the other as an uninfected control, and sampled at the time points indicated in **Supplementary Figure 2B**. A sample just prior to T2 addition served as the T0 time point. For the T0, 3 ml of culture was collected, whilst subsequent time points, 2 ml of sample. All samples for **Figure 3** and **Supplementary Figure 2B** were OD_600_ synchronised, pelleted, resuspended in 100 μl of x2 SDS-loading buffer (Sigma) and 10 μl was separated on a 4-15% gradient denaturing gel (Biorad). Immunoblotting was conducted in accordance with standard laboratory protocols, with primary antibodies incubated overnight. Commercially available primary mouse monoclonal ɑ-His (Sigma, SAB2702219), ɑ-3XFLAG (Sigma, F1804), and ɑ-DnaK (Enzo, 8E2/2) antibodies were used at 1:5000 dilution, whilst commercially available secondary antibody HRP ɑ-mouse IgG (BioLegend, 405306) was used at 1:10000 dilution. ECL Primer western blotting detection reagent (GE Healthcare, RPN2232) was used to develop the blots, which were analysed on the ChemiDoc MP imaging system, and bands quantified using Image Lab software.

## Results

### T2 requires Hfq for optimal replication in *E. coli*

Hfq has previously been implicated in multiple aspects of phage–host interaction, including various phages infecting *E. coli* (10). To determine whether Hfq is required by T2 for post-transcriptional regulation, we amplified T2 in WT, Δ*hfq* and trans-complemented (Δ*hfq*::*hfq*) bacteria, serially diluted the lysates, spotted onto a lawn of WT *E. coli* and enumerated the number of infecting particles (phage titre) in each lysate by counting plaque forming units (PFU). In Δ*hfq*::*hfq* bacteria, Hfq was plasmid-borne (pHfq) and induced upon addition of L-arabinose. As shown in **Figure 1A**, the titre of T2 (PFU/ml) from Δ*hfq* bacteria was decreased ∼11 fold compared to T2 titre from WT bacteria, suggesting that T2 requires Hfq to optimally replicate in *E. coli*. Consistent with this view, the T2 titre from Δ*hfq*::*hfq* bacteria was increased by ∼30 fold compared to that from WT bacteria, suggesting that overexpression of Hfq markedly benefits T2 replication. Hfq has pleiotropic functions in bacterial physiology and several residues at all three surfaces of its conserved toroidal core contribute to its diverse functions. Conserved amino acids residues Q8, D9, R16, R17 and Y25 have been widely implicated in post-transcriptional regulation (19,20). Therefore, we next measured T2 titres in lysates from Δ*hfq*::*hfq* bacteria expressing Hfq mutant variants harbouring an alanine substitution at Q8, D9, R16, R17 and Y25. As shown in **Figure 1A**, the T2 titres from Δ*hfq*::*hfq* bacteria expressing the Hfq mutants were reduced by ∼4 to ∼18 fold compared to that from WT bacteria. Further, the carboxyl-terminal tail of Hfq acts synergistically with the other RNA-binding faces on the conserved core and contributes to the specificity of its RNA annealing activity. However, the absence of the carboxyl-terminal tail of Hfq (Δ*hfq* + pHfq ΔCTD) did not significantly affect T2 replication (**Figure 1A**; also see below). Overall, we conclude that Hfq-mediated post-transcriptional regulation is a requirement for optimal T2 replication in *E. coli*.

**Figure 1.**
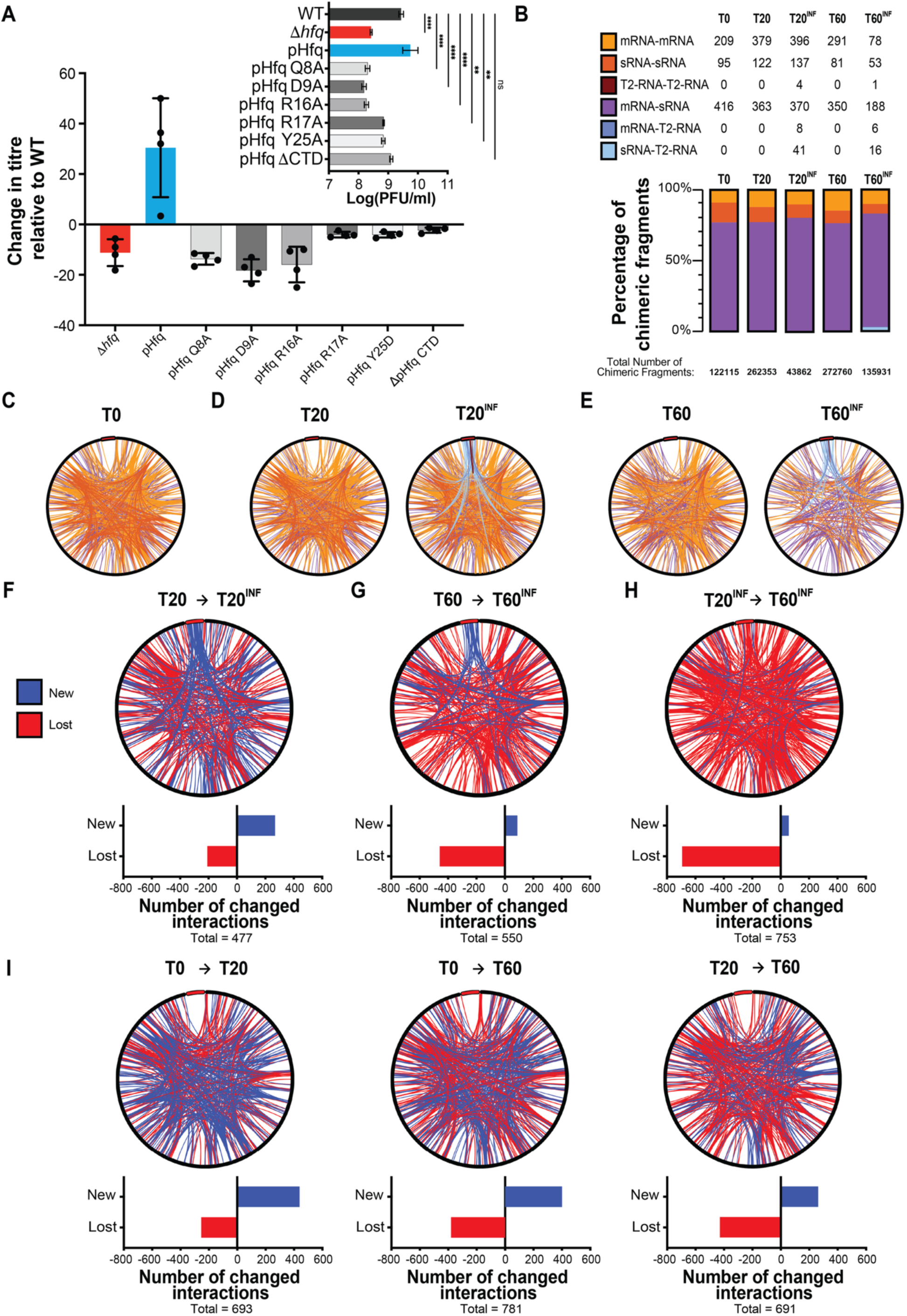
RIL-seq reveals reprogramming of Hfq-mediated RNA interactome during T2 replication. (**A**) Graph showing the fold-change in phage titre relative to WT in bacteria which are either lacking Hfq or contain point mutants of Hfq. Inset shows phage titre as PFU/ml used to calculate fold-change. Error bars represent standard error (n= 4). Statistical analysis was performed by One-Way ANOVA (****P< 0.0001; **P< 0.01). (**B**) Bar chart showing the relative frequencies of each RNA-RNA interaction type found in chimeric fragments, colour-coded as in the table which shows the number of unique chimeras present in the two RIL-seq data sets. (**C**) Circos plot of RNA-RNA interactions represented by at least 10 chimeric fragments, mapped to the *E. coli* MG1655 and T2 phage genomes at T0 (i.e. before T2 infection). The T2 genome is indicated as a red bar at the 12 o’clock position. Different types of RNA-RNA interactions are colour-coded as in (B). (**D**) As in (C) but for 20 min after T2 infection (T20^INF^). The corresponding uninfected control is indicated as T20. (**E**) As in (C) but for 60 min after T2 infection (T60^INF^). The corresponding uninfected control is indicated as T60. (**F-H**) Circos plots representing the newly established (blue) and lost (red) RNA-RNA interactions between the indicated time points. The graphs below indicate the numbers of newly established and lost interactions for each time point comparison; the total number of changed interactions is also given. (**I**) As in (F-H) but for the uninfected control samples.

### T2 infection causes global transcriptome changes

To better understand the requirement for Hfq for optimal T2 replication, we initially determined the global transcriptional changes that occur in T2 infected *E. coli* by RNA-seq. We infected exponentially-growing (OD_600_ of ∼0.3) *E. coli* with T2 (at a multiplicity of infection of 20) and sampled at 20 (T20^INF^) and 60 (T60^INF^) minutes post-infection. In parallel, we sampled from an uninfected culture at T20 and T60. A sample taken immediately prior to the addition of the phage to the bacterial culture served as the T0 time point (see later). We observed that bacteria continued to replicate following infection, but the growth of the infected culture became attenuated ∼90 min after infection and an obvious culture collapse (indicative of phage-mediated cell lysis) was not detected even by 4 h after infection (**Supplementary Figure 1A**). However, enumeration of the proportion of viable cells by counting colony forming units (CFU) in the infected culture showed a dramatic decrease ∼60 min after infection, suggesting successful adsorption and replication of T2 (**Supplementary Figure 1B**). Indeed, a clear collapse of the infected culture was seen eventually, ∼5 h after infection (**Supplementary Figure 1C**). Comparison of the transcriptomes of uninfected and infected time points revealed, unsurprisingly, upregulation of T2 transcripts (defined as those transcripts with expression levels changed ³ 2-fold with a false discovery rate adjusted p≤0.05). As shown in **Supplementary Figure 1D and 1E**, we observed that 31.6% of the T2 transcriptome was upregulated at T20^INF^ and 91.0% at T60^INF^. We further observed 32 and 44 downregulated and 225 and 277 upregulated *E. coli* genes at T20^INF^ and T60^INF^, respectively, showing reprogramming of host gene expression during T2 replication in *E. coli*. Most of the differentially expressed genes at T20^INF^ were associated with carbohydrate transport and metabolism; however, at T60^INF^, differentially expressed genes were associated with diverse cellular functions (**Supplementary Figure 1F**). Notably, of the >100 known *E. coli* non-coding RNAs (including sRNAs) (21), 5 were differentially expressed (all upregulated) at T20^INF^, whilst 15 were differentially expressed (6 upregulated; 9 downregulated) at T60^INF^ (**Supplementary Figure 1G and 1H**). We note that, at T20^INF^, although the functions of many upregulated sRNAs are not well-defined, the upregulation of SgrS, is consistent with the perturbed expression of genes associated with carbohydrate transport and metabolism (e.g. *ptsG* (22), *manXYZ* (23), *yigL* (24)) (**Supplementary Figure 1F**). Similarly, at T60^INF^, many sRNAs that globally affect gene expression (*e.g.* SdsR (13), DsrA (25)) were downregulated, consistent with the perturbated expression of genes across diverse cellular functions. We note that at T60^INF^, RyfA, a sRNA that does not rely on Hfq but on the RNA chaperone ProQ (26), is upregulated, possibly suggesting an involvement of ProQ as well as Hfq during T2 replication. We further note that non-coding RNA SsrS is upregulated at T60^INF^. SsrS encodes the 6S RNA which binds to the housekeeping RNAP form containing the σ^70^ promoter-specificity factor and acts to reduce its activity during stress (e.g. during transition from exponential to stationary phase of growth). By doing so, it causes the dissociation of σ^70^ from the RNAP and enables it to bind to σ^S^, the central stress response promoter-specify factor and thus adapts gene expression to stress (27,28). The observation that SsrS becomes upregulated at T60^INF^ might suggest a host response to stress imposed by T2 replication. Although our results indicate the involvement of several sRNA during T2 replication, the expression levels of the majority of Hfq-dependent *E. coli* sRNAs did not change during T2 replication. This observation could potentially underscore the need for host sRNAs for optimal T2 replication, consistent with the requirement of Hfq (**Figure 1A**).

### Extensive reprogramming of the Hfq-mediated RNA-interactome during T2 replication

Hfq-associated sRNAs commonly act through base-pairing with other transcripts to modulate translation and RNA stability (29). Thus, we used RIL-seq (RNA interaction by ligation and sequencing) to define the Hfq-mediated RNA-RNA interactions (hereafter simply RNA-RNA interactions) that occur during T2 replication at the transcriptome-wide scale (30). RIL-seq allows isolation and sequencing of ligated RNA–RNA pairs (chimeras) through immunoprecipitation of Hfq-RNA complexes. To immunoprecipitate Hfq-associated RNA-RNA interactions during T2 replication, we used an *E. coli* strain containing a 3XFLAG-tag sequence fused C-terminally to *hfq* at its native chromosomal location (13). All RNA-RNA pairs that were present in at least one of the three biological replicates (note, T20^INF^ only consisted of two biological replicates) but were represented by at least 10 chimeric fragments were considered for downstream analysis. This gave a total of 122115, 262353, 43862, 272760, 135931 chimeric fragments at T0, T20, T20^INF^, T60 and T60^INF^, respectively (**Figure 1B**). The total number of chimeric fragments captured at each timepoint is large. This is due to the stringency of our cut-off of 10, which ensured that the relatively low abundances of chimeric fragments containing phage transcripts (hereafter T2-RNA) were captured. At all the time points, the largest proportion of chimeric fragments consisted of a bacterial sRNA and a mRNA. Hereafter all bacterial mRNA and bacterial sRNA are simply referred to as mRNA and sRNA, respectively. Of the total number of chimeras detected at each point, the numbers of unique chimeras of different types are summarised in **Figure 1B**. Consistent with previous Hfq-mediated RNA interactomes (hereafter simply RNA interactomes), we detected numerous sRNA-mRNA and sRNA-sRNA chimeras. Of the 15 differentially expressed sRNAs across T20^INF^ and T60^INF^ (**Supplementary Figure 1G and 1H**), only 5 sRNAs (CsrB, DsrA, FnrS, MicF and SdsR) that are downregulated at T60^INF^ formed chimeric fragments associated with Hfq. In the infected samples, ∼0.1-2% of the total number of chimeric fragments consisted of either a sRNA or mRNA and a T2-RNA. At T20^INF^ and T60^INF^, we detected several RNA-RNA interactions between a sRNA and a T2-RNA (41 and 16, respectively), mRNA and a T2-RNA (8 and 6, respectively) and between two T2-RNA (4 and 1, respectively) (**Figure 1B**). We generated circos plots in which unique RNA-RNA interactions represented by at least 10 chimeric fragments mapped to the *E. coli* and T2 genomes. In the circos plots the T2 genome was ‘integrated’ at the 12 o’clock position. Visual comparison of the circos plots showing all RNA-RNA interactions at T0 (**Figure 1C**), T20 (**Figure 1D**) and T60 (**Figure 1E**) in uninfected cells revealed little detectable differences. However, comparison of corresponding circos plots representing the unique RNA-RNA interactions in the infected cells revealed a clear ‘reprogramming’ of the Hfq-mediated RNA-interactome during T2 replication (**Figure 1D** and **1E**). Indeed, at T20^INF^ ∼268 new Hfq-mediated RNA-RNA interactions are established, whilst ∼209 RNA-RNA interactions are lost when compared to T20 (**Figure 1F**). At T60^INF^, ∼90 new RNA-RNA interactions are established but a greater number, ∼460, of RNA-RNA interactions are lost (**Figure 1G**). Therefore, between T20^INF^ and T60^INF^, the majority of RNA-RNA interactions are lost, with few (∼60) new ones established, leading to a collapse of the Hfq-associated RNA-interactome in the T2 infected cells (**Figure 1H**). In other words, ∼63% of RNA-RNA interactions that existed at T0 were lost by T60^INF^. In contrast, although the RNA-interactome was dynamic in the T2 uninfected sample during the time course of our experiment (*i.e.* 60 min), a total collapse of the RNA-interactome was not evident (**Figure 1I**). This apparent reprogramming of the RNA-interactome during T2 replication was not due to gross changes in total RNA concentration between the T2 infected and uninfected cells (**Supplementary Figure 2A**) or differences in Hfq protein levels (**Supplementary Figure 2B**). Further, we identified several bacterial RNA-RNA interactions (121 mRNA-sRNA, 34 sRNA-sRNA and 49 mRNA-mRNA) that remained unaffected during T2 replication (**Supplementary Figure 3**). In other words, ∼23% of the RNA-RNA interactions that existed at T0 were retained at T60^INF^, suggesting that the regulatory consequences of these interactions (hereafter core interactions) might be essential to support cellular functions during T2 replication. We note that 3 (DsrA, FnrS, MicF) of the 5 sRNAs that were downregulated at T60^INF^ are involved in core interactions. The relative abundance of chimeric fragments representing each of the core RNA-RNA interaction types differed between T0 and T60^INF^ (**Supplementary Figure 4**), which is further supportive of the view that the Hfq-mediated RNA interactome is reprogrammed during T2 replication. Overall, the results reveal that the RNA-interactome is extensive and dynamic during T2 replication in *E. coli* but eventually becomes destabilised, indicative of phage takeover of the bacterial cell.

### T2 co-opts the bacterial sRNA ArcZ for optimal replication in *E. coli*

Phage infection typically relies on several host factors for adsorption, replication, and assembly (31). To understand whether host-derived sRNAs are required during T2 replication, we focused on the RNA-RNA interactions involving a T2-RNA. We detected several unique T2-RNA-sRNA (57), T2-RNA-mRNA (15) and T2-RNA-T2-RNA (5) interactions (**Supplementary Figure 5**). The majority of the interactions involving a T2-RNA occurred with sRNAs, and at T60^INF^ most of T2-RNA-sRNA interactions involved the sRNA ArcZ (**Figure 2A**), a major post-transcriptional regulator in *E. coli* and related species (32–36). ArcZ interacts with five protein-coding T2 transcripts, which correspond to the T2 genes *arn.2* (unknown protein; 11 kDa), *tk.4* (putative thymidine kinase; 17 kDa), *soc* (putative capsid protein; 9 kDa), *23* (putative capsid protein; 56 kDa) and *55* (putative promoter specificity factor; 22 kDa) (**Figure 2B**). To confirm that ArcZ is required for optimal T2 replication, we compared T2 titres in lysates from WT, Δ*arcZ* and Δ*arcZ*::*arcZ* bacteria. In Δ*arcZ*::*arcZ* bacteria, ArcZ was plasmid-borne (pArcZ) and expressed upon induction with IPTG. As shown in **Figure 2C**, we observed a ∼2.5-fold reduction in T2 titre in the lysates from Δ*arcZ* compared to WT lysate. As expected, T2 titre was restored to near WT level in the lysates from Δ*arcZ*::*arcZ* bacteria. We then compared the proteomes of WT and Δ*arcZ* bacteria at T60^INF^, focusing particularly on T2 proteins, to understand ArcZ’s influence on both the expression of its targets captured by RIL-seq and global T2 gene expression. We identified 43 T2 proteins, representing ∼19.5% of the T2 proteome. This included 3 of the 5 gene products of putative ArcZ targets captured by RIL-seq (*tk.4*, *23* and *55*) (**Figure 2B**). However, products of the other T2 transcripts targeted by ArcZ based on our RIL-seq analysis, *arn.2* and *soc* were not present in the proteome data, as their molecular weights are below the detection threshold. As shown in **Figure 2D**, none of the detected T2 proteins were differentially expressed in Δ*arcZ* bacteria, suggesting that the interaction between ArcZ and the *tk.4*, *23* and *55* T2 transcripts might not be productive or involve a different mechanism of regulation (37). As shown in **Figure 2E**, *tk.4* and *23* T2 transcripts are also targeted by other sRNA and the *23* T2 transcript also interact with another T2-RNA (22). Therefore, their regulatory fate might not be solely dependent on ArcZ. On the host side, as shown in **Figure 2D**, one protein was downregulated and 13 proteins were upregulated in Δ*arcZ* bacteria. One of the downregulated proteins was σ^S^, whose translation is dependent on three different sRNAs, including ArcZ (33,38). To rule out that the replication deficiency of T2 in Δ*arcZ* bacteria was not due to compromised σ^S^ availability, we compared T2 titres from WT, Δ*arcZ* and Δ*rpoS* bacteria. As shown in **Figure 2F**, the T2 titres from WT and Δ*rpoS* bacteria were comparable, suggesting that the replication deficiency of T2 in Δ*arcZ* bacteria is due to the absence of ArcZ and not due to the compromised σ^S^ availability. As the highest proportion of ArcZ chimeras involved the *arn.2* transcript (**Figure 2B**), the *arn.2* transcript was solely targeted by ArcZ (**Figure 2E**) and detected in libraries of all three biological replicates (**Supplementary Data 3**), we focused on the regulation of *arn.2* gene expression by ArcZ and its relevance to T2 replication. The *arn.2* gene is part of a 5 gene transcript (hereafter *arn* transcript) and represents one of the first middle gene clusters on the T2 genome. The *arn* transcript consists of *arn.4*, *arn.3*, *arn.2*, *arn.1* and *arn,* from 5’ to 3’. As shown in **Supplementary Figure 6**, RNA corresponding to *arn.4*, *arn.3*, *arn.2*, *arn.1* and *arn* is detected in high abundance in the transcriptomes of T60^INF^. The functions of proteins encoded by *arn.4*, *arn.3*, *arn.2*, *arn.1* are not known (see discussion). However, the product of *arn*, Arn (anti restriction nuclease) is an inhibitor of McrBC restriction endonuclease, which functions as a general defence mechanism to specifically degrade any invading foreign DNA (*e.g.* that of T2) containing 5-hydroxymethylcytosine (39). As the absence of ArcZ compromises T2 replication, we posited that ArcZ positively influences the *arn* transcript, including the expression of Arn. Thus, we considered that the T2 replication deficiency in Δ*arcZ* bacteria may be rescued by deleting the *mcrC* gene (which abrogates McrBC activity). As shown in **Figure 2G**, T2 titres from WT and Δ*arcZ*Δ*mcrC* bacteria were comparable, supporting the view that ArcZ positively influences the *arn* transcript and consequently Arn expression. We conclude that ArcZ is co-opted by T2 to facilitate the expression of a protein required for countering a host defence mechanism, allowing optimal replication in *E. coli*. We also note that the stability of ArcZ is not affected by the CTD of Hfq (40), which is consistent with our results that T2 titres from Δ*hfq* + pHfqΔCTD bacteria is comparable to those in WT bacteria (**Figure 1A**). Further, although the functions of *arn.4*, *arn.3*, *arn.2* and *arn.1* remain unknown, it is conceivable that they too are involved in counter-defence functions.

**Figure 2.**
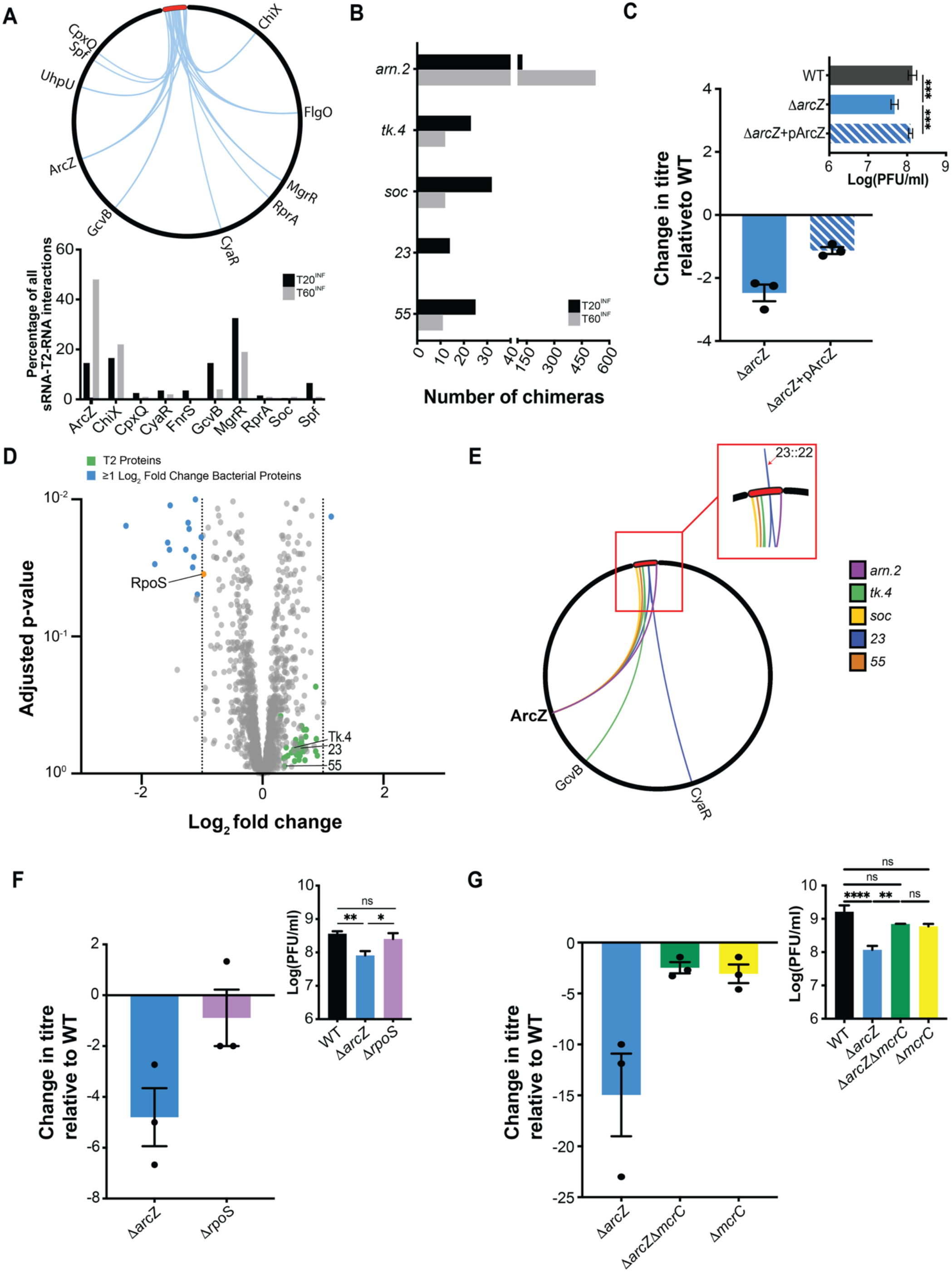
RIL-seq reveals a requirement of the host nucleic acid factor ArcZ for optimal replication of T2 phage in *E. coli*. (**A**) Circos plot representing all interactions between sRNA and T2-RNA (see Supplementary Figure 5 for details). The graph below shows sRNA-T2-RNA interactions as a percentage of all sRNA-T2-RNA interactions occurring at either T20^INF^ and T60^INF^. (**B**) Graph representing the number of chimeras detected for each interaction involving ArcZ and a T2-RNA at either T20^INF^ and T60^INF^. (**C**) Graph showing the fold-change in phage titre relative to WT in both Δ*arcZ* and Δ*arcZ* bacteria complemented with pArcZ. Inset shows phage titre as PFU/ml used to calculate fold-change. (**D**) Volcano plot of differential protein levels at T60^INF^ in Δ*arcZ* bacteria compared to WT bacteria shown as a log_2_ fold change. Proteins were considered differentially expressed (defined as those proteins with expression levels changed ³ 2-fold with a false discovery rate adjusted p≤0.05) and indicated in blue. Phage proteins are indicated in green. Bacterial proteins whose abundance did not change are in grey. T2-RNA targets of ArcZ captured by RIL-seq are labelled; σ^S^ (RpoS) is shown in orange. Bacterial proteins whose abundance did not change significantly are shown in grey. (**E**) Circos plot representing T2-RNA which interact with ArcZ and any of their additional interactions with other sRNA and/or T2-RNA. The T2 genome is amplified in the inset to show T2 transcript *23* interacts with T2 transcript *22*. (**F**) Graph showing the fold-change in phage titre relative to WT in both Δ*arcZ* and Δ*rpoS* bacteria. Inset shows phage titre as PFU/ml used to calculate fold-change. (**G**) Graph showing the fold-change in phage titre in Δ*arcZ*, Δ*arcZ*Δ*mcrC* and Δ*mcrC* bacteria relative to WT. Inset shows phage titre as PFU/ml used to calculate fold-change. In (C), (F) and (G) error bars represent standard error (n=3). Statistical analysis was performed by One-Way ANOVA (****P< 0.0001; ***P<0.001 and **P< 0.01).

### ArcZ prevents RNase E-mediated degradation of the *arn* transcript

The *arn* operon is found in diverse lytic phages (**Supplementary Figure 7**) (41). The predicted binding region of ArcZ lies within the 297 nucleotides of *arn.2* and base-pairing interactions between ArcZ and *arn.2* occurs at the *arn.2* 3′ end between positions +285 and +294, relative to its translation start site (**Figure 3A**). Indeed, both this ArcZ binding region and ArcZ itself are conserved within diverse T2-like phages that contain a homologous *arn* operon (**Supplementary Figure 7**) and their respective host bacteria. Closer inspection of the *arn* operon revealed that the ArcZ binding region directly precedes an intergenic region (IR) 70 nucleotides in length, at the end of which we identified a putative RNase E binding region (5’-GcAUU-3’)- located 11 nucleotides upstream of the *arn.1* translation start codon (**Figure 3A**). The RNase E binding region is also conserved in most phages that contain the *arn* operon (**Supplementary Figure 7**). To understand the mechanism by which ArcZ positively influences the *arn* transcript, we developed a minimal reporter assay. For this, we made a plasmid construct, hereafter called construct I, which contained the DNA sequence of both *arn.2* (containing the ArcZ binding region) and the IR (containing the RNase E binding region), with an in-frame fusion of a 6xHis-tag at the 5’-terminal end of *arn.2*, placed downstream of a L-arabinose inducible promoter. This allowed us to detect 6xHis-Arn.2 (hereafter simply Arn.2) in whole-cell extracts of bacteria by immunoblotting with an ɑ-His antibody to evaluate the influence of ArcZ and RNase E on Arn.2 production. As shown in **Figure 3B**, upon induction in WT bacteria, we were able to detect Arn.2 expression. In contrast, the Arn.2 protein was not detected in whole-cell extracts of Δ*arcZ* bacteria, supporting the view that ArcZ positively impacts the transcript from construct I (*i.e.* 6xHis-*arn.2*-IR). We were unable to recover Arn.2 expression when pArcZ was provided to Δ*arcZ* bacteria for reasons we do not understand (but see below). The removal of the IR from construct I, generating construct II, resulted in improved expression of Arn.2 in WT bacteria. Notably, expression of Arn.2 in Δ*arcZ* bacteria was comparable to that seen in WT bacteria (**Figure 3C**). In other words, Arn.2 expression from construct II, when the entire IR including the RNase E binding region, was absent, became ArcZ independent. Construct III, in which the IR was shortened from the 3’ end by 20 nucleotides to remove the putative RNase E binding region, revealed comparable levels of Arn.2 detected in whole-cell extracts of both WT and Δ*arcZ* bacteria (**Figure 3D**). We thus posited that the binding of ArcZ might protect the *arn* transcript from RNase E-mediated degradation in *E. coli*, despite the putative RNase E and ArcZ binding regions being located 57 nucleotides from each other. The results suggested that the *arn* transcript is degraded by RNase E when ArcZ is absent. As RNase E primarily exhibits endonuclease activity in the 5’ to 3’ direction by recognising a 5’ monophosphate (42,43), it is likely that the 5’-3’ product that results from the cleavage of the transcript originating from construct I is degraded by other cellular 3’ to 5’ exonucleases (*e.g.* by RNase E associated PNPase), thus leading to absence of His-Arn.2 expression in Δ*arcZ* bacteria (**Figure 3B**). However, we also note that RNase E also exhibits 3’ to 5’ cleavage directionality (44). As only Arn (the inhibitor of McrBC restriction endonuclease), which encoded by the last gene on the *arn* operon, is functionally characterised, we wanted to better connect the proposed mechanism by which ArcZ influences the *arn* transcript with the data shown in **Figure 2G** (*i.e.* the suppression of T2 replication deficiency in a Δ*arcZΔmcrC E. coli* strain). For this, we generated construct IV, in which the DNA sequence corresponding to the entire *arn* operon was placed downstream of a L-arabinose inducible promoter with an in-frame fusion of a 6xHis-tag at the 3’-terminal end of *arn.* This allowed us to monitor expression of Arn-His and also served as a proxy for RNase E-mediated degradation of the *arn* transcript in the 5’ to 3’ direction. As shown in **Figure 3E**, we detected gradual accumulation of Arn in WT bacteria following induction; in contrast, in Δ*arcZ* bacteria, Arn-His accumulated markedly slower following induction. As a control experiment, we used Δ*hfq* bacteria. Here, any effect of ArcZ on Arn-His expression should be abolished as Hfq is required for both processing of ArcZ (32–34,45) and facilitating base pairing interactions between ArcZ and the *arn* transcript. As shown in **Figure 3E** (bottom panel), and as expected, we did not detect Arn-His expression in Δ*hfq* bacteria but Arn-His expression was recovered when Δ*hfq* bacteria containing pHfq. We next compared T2 titres from WT and Δ*arcZ* bacteria in which the whole *arn* transcript was overexpressed (from construct VI). As shown in **Figure 3F**, titres of T2 from WT bacteria were similar regardless of whether the *arn* operon was overexpressed. However, overexpression of the *arn* operon did not revert T2 titre from Δ*arcZ* bacteria to levels seen in WT lysates, affirming the positive influence of ArcZ on the *arn* transcript. We thus posited that overexpression of the *arn* operon in which the RNase E binding region in the IR is mutated, *i.e.* changed from 5’-GcAUU-3’ to 5’-GcAGG-3’ (thereby preventing RNase E-mediated cleavage and degradation (45); construct V) would revert T2 titres from Δ*arcZ* bacteria to levels seen in WT bacteria. As shown in **Figure 3F**, this was indeed the case. We note T2 titres from WT bacteria were similar regardless of whether the *arn* operon from construct V was overexpressed. We conclude that ArcZ functions to prevent RNase E-mediated degradation of the *arn* transcript to counteract a host antiviral defence system.

**Figure 3.**
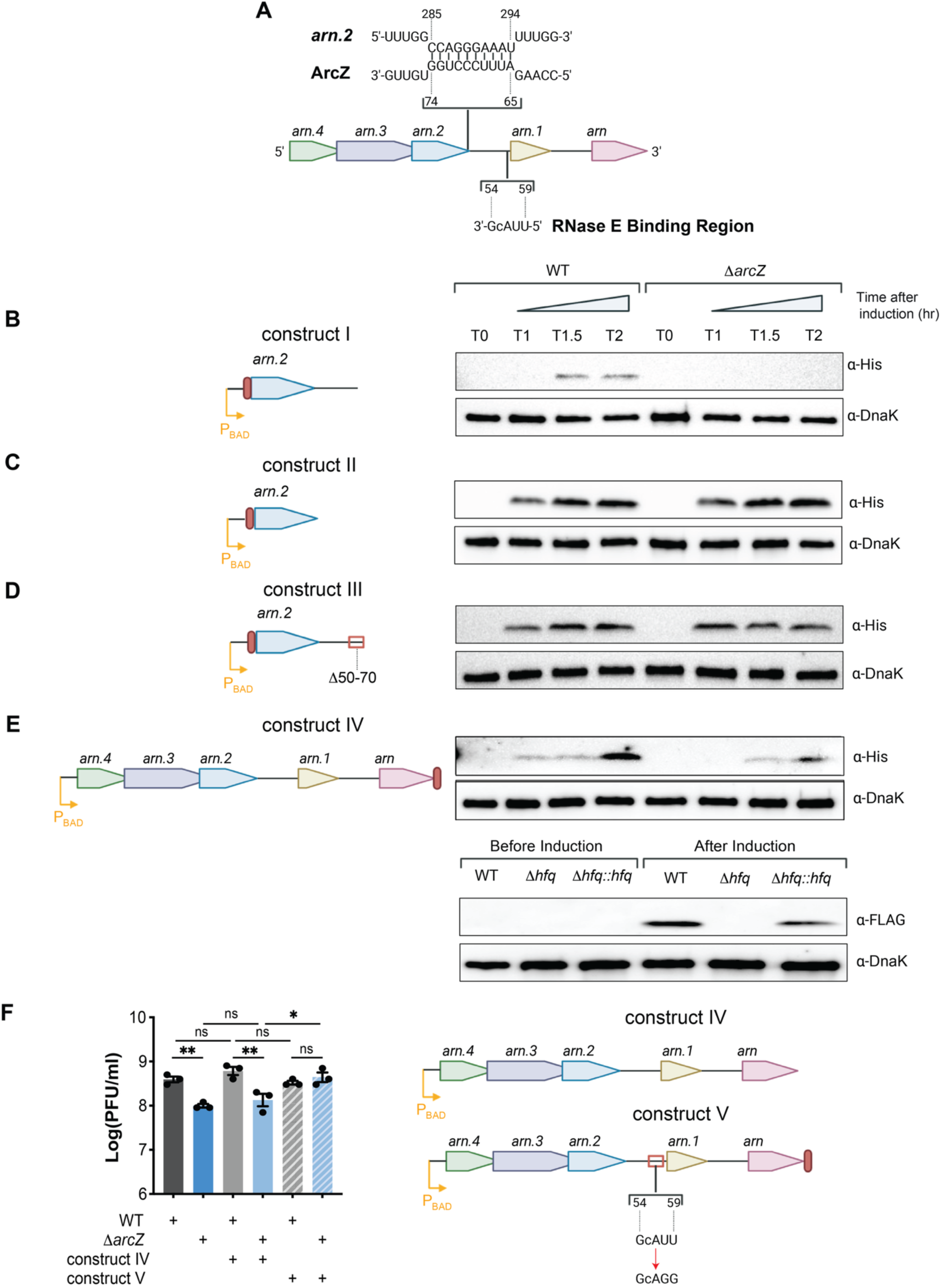
ArcZ prevents RNase E-mediated degradation of the *arn* transcript during T2 replication in *E. coli*. (**A**) Schematic representation of the *arn* transcript, ArcZ and RNase E binding regions are indicated. (**B-E**) Representative immunoblots of whole cell extracts of *E. coli*. The strain background, induction time points and plasmid constructs are indicated (see text); T0 indicates the time point prior to L-arabinose addition. Immunoblots were probed with ɑ-His antibody to detect each 6xHis-tagged protein and ɑ-DnaK antibody to detect DnaK as the loading control. Each construct is schematically represented on the left of the corresponding immunoblot with the red bar illustrating the 6xHis-tag position. Experiments using Δ*hfq* bacteria were induced for 2 h. (**F**) Graph representing phage titres in WT and Δ*arcZ* strains in which either the whole arn operon (construct IV) or the whole *arn* operon with the RNase E binding region mutated (construct V) was overexpressed. The error bars represent standard error (n=3). Statistical analysis was performed by One-Way ANOVA (**P< 0.01; ns=not significant).

## Discussion

Phages continuously evolve strategies to hijack host processes to optimise their own replication (46). Previous work has shown how phages produce Hfq-dependent sRNA to subvert bacterial processes (11,12). Here we uncover a previously unrecognised regulatory layer in phage-bacteria interactions in which a Hfq-dependent bacterial sRNA is required to meet a phage gene expression need. Using T2 infection of *E. coli* as a model, we show that the sRNA ArcZ, a ubiquitous post-transcriptional regulator in Enterobacterales (32), is co-opted by T2 to stabilise a phage transcript (the *arn* operon) encoding a protein, amongst others, that antagonises a bacterial restriction–modification defence system (McrBC). Our finding expands the molecular armament in the phage–bacteria conflict by revealing that phages can exploit not only host proteins but also host nucleic acid factors for their own needs. The discovery that ArcZ enhances T2 replication by protecting the *arn* transcript from RNase E degradation represents a new role for bacterial sRNAs in phage biology. To the best of our knowledge, the idea that phages could co-opt bacterial sRNA to post-transcriptionally regulate their gene expression has not been documented. Several phages encode proteins that inhibit RNase E directly or redirect its activity (5,6,8), but the use of ArcZ by T2 phage provides an alternative, RNA-mediated route to the same end, which is to prevent the degradation of specific phage transcripts without encoding a dedicated RNase E inhibitor. Indeed, we did not detect homologues of known phage-encoded RNase E modulators (e.g. *E. coli* phage T7 Gp0.7 or *Pseudomonas aeruginosa* phage ϕKZ Dip) in T2. This reinforces the view that phages can also tap into pre-existing bacterial gene regulatory circuitry instead of evolving large repertoires of viral regulators, allowing their genomes to be compact and efficient. Additionally, by protecting the *arn* transcript, ArcZ indirectly mitigates bacterial McrBC restriction activity and facilitates efficient phage DNA replication. This mechanism differs fundamentally from previously described phage tactics involving Hfq and regulatory RNA molecules: RNA phages such as Qβ recruit Hfq as an unwinding chaperone for viral RNA replication (9), and several DNA phages encode their own Hfq-dependent sRNAs that sponge or modulate host transcripts (11,12). In contrast, our study shows that T2 phage appropriates a bacterial sRNA to stabilise a key counter-defence mechanism to evade the bacterial McrBC defence mechanism. The observation that deletion of *arcZ* compromises T2 replication, whereas loss of *mcrC* suppresses this defect, firmly connects ArcZ activity to the restriction-evasion pathway by T2 phage, via post-transcriptional regulation of the *arn* transcript. Except for Arn, the inhibitor of McrBC, the functions of the other proteins encoded by the *arn* transcript are unknown, yet it is conceivable that they also have functions in counter-defence.

The RIL-seq data revealed a striking collapse and reorganisation of the Hfq-mediated RNA interactome during T2 replication. Early in infection, several new RNA–RNA interactions emerge, including interactions between bacterial sRNAs and phage transcripts. As infection progresses, the global RNA-RNA interaction network progressively disassembles. This highlights the broader domination of the bacterial cell by T2. Whether this destabilisation during T2 replication results from competition for Hfq (11,34,47,48), or from stress-induced changes to cellular physiology, remains unknown and requires further investigation. As the relative abundance of non-coding RNAs (including sRNAs) do not change during T2 replication, as revealed by the RNA-seq data, it is unlikely that the collapse of the Hfq-mediated RNA-interactome is due to destabilisation of sRNAs. However, as exemplified by ArcZ–*arn.2* interaction, our results indicate a fine-tuned modulation rather than total inhibition of host RNA metabolism during the course of T2 replication. Approximately one quarter of Hfq-mediated interactions between bacterial RNAs persist during T2 replication. This may represent a “core” post-transcriptional network that needs to be maintained to ensure host survival for as long as possible until T2 replication is complete. Further, we identified 4 unique interactions between two T2 transcripts and 11 unique interactions between T2 transcripts and bacterial mRNA (**Supplementary Figure 3**) during T2 replication. It is possible that the former represents an example of how T2 appropriates Hfq and the latter represents an example of how T2 modulates bacterial genes expression using phage sRNA. Both strategies could serve to optimise development of T2 progeny in *E. coli*.

The *arn* operon is widespread among lytic phages infecting Enterobacteriaceae (41), and both the ArcZ and RNase E binding regions are conserved (**Supplementary Figure 7**), implying evolutionary pressure to maintain host-dependent regulatory control and could represent a mechanism of host tropism. This conservation suggests a generic strategy wherein diverse phages exploit the conserved Hfq-dependent sRNA ArcZ to stabilise the *arn* operon. Although, the involvement of other sRNA in *arn* transcript regulation in other phages cannot be excluded. In the case of T2, from the host perspective, the co-option of ArcZ could impose a selection pressure for modifying ArcZ binding properties or for deploying competing sRNA to sponge ArcZ. However, given that ArcZ’s interactions with bacterial transcripts during T2 replication is extensive (**Supplementary Figure 8**), it is unlikely that a strategy to modulate ArcZ’s activity would be viable. Nonetheless, our discovery broadens our understanding of the evolutionary arms race between phages and bacteria that occur at the post-transcriptional level.

In sum, our study provides the first evidence that a bacterial sRNA directly promotes phage gene expression. By stabilising the *arn* transcript, ArcZ ensures the production of a counter-defence protein(s) essential for efficient T2 replication. We showed that ArcZ prevents RNase E-mediated degradation of the *arn* transcript, a regulatory mechanism that has documented for other sRNAs as well (24,49). The ArcZ and the RNase E binding region are separated by several nucleotides, implying either a conformational rearrangement upon ArcZ binding that sequesters the RNase E cleavage motif, or recruitment of Hfq to remodel the local RNA architecture to prevent RNase E binding. Thus, future work will need to focus on the biochemical aspects of this interaction to establish the precise mechanism of action of the ArcZ-Hfq-RNase E axis on the regulation of the *arn* transcript. The conceptual advance in our understanding of phage-bacteria interaction uncovered in this study, that phages harness endogenous regulatory RNA molecules to fine-tune their own gene expression, clearly extends beyond the T2-*E. coli* system. This offers new insight into the interplay and co-evolution of bacterial post-transcriptional networks and the viruses that invade them. Notably, the discovery of phages that require host nucleic acid factors for optimal replication, particularly for expressing counter-defence proteins, could be leveraged to bioengineer synthetic therapeutic phages that escape target bacterial counter-defence systems.

## Data availability

The RNA-seq and RIL-seq data discussed in this publication are accessible through Gene Expression Omnibus (GEO) under the following accession codes: GSE330885 (RNA-seq) and GSE331271 (RIL-seq). The proteomics data can be accessed through PRIDE: PXD078616.

## Acknowledgements

We thank members of the S.W. and K.P. laboratories for constructive comments and discussion on this work.

## Funding

Work in S.W. laboratory was funded by grants by the Biotechnology and Biological Sciences Research Council Research Grant [BB/Y009630/1] and Leverhulme Trust Research Project Grant [RPG-2025-105]. Work in K.P. laboratory was funded by the German Research Foundation (PA2820/3-2 – Project number 464459808 and EXC 2051 – Project number 390713860) and the European Research Council (ArtRNA, CoG-101088027). Funding for open access charge was obtained via the Imperial College UKRI OA fund.

## Notes

### Competing Interest Statement

The authors have declared no competing interest.

