## Supplementary Figures 1-8 for "A nucleic acid host factor enables optimal phage replication in *Escherichia coli*"

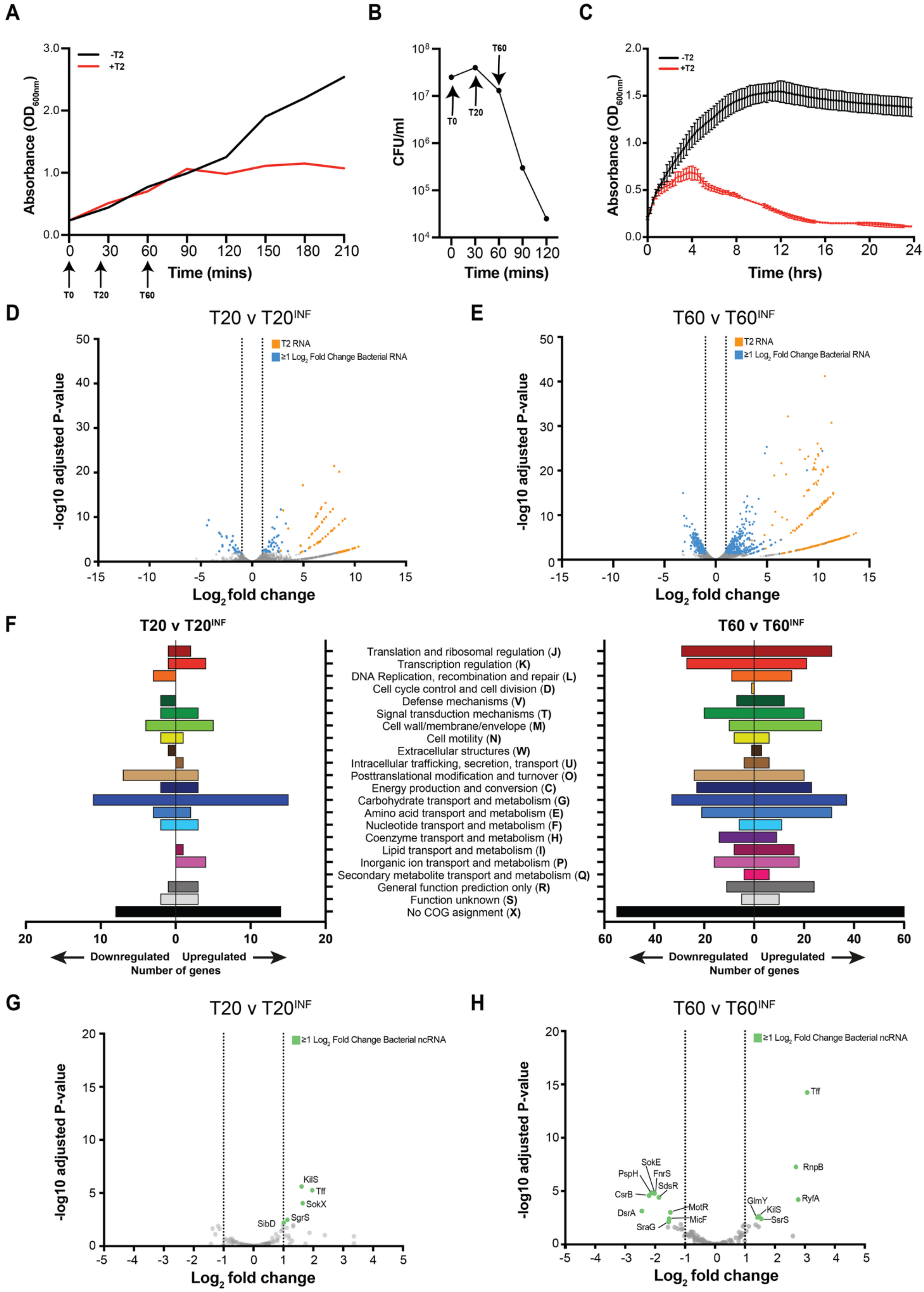
A
B
C
D
E
F
G
H
T20 v T20INF
T60 v T60INF
T2 RNA
≥1 Log2 Fold Change Bacterial RNA
≥1 Log2 Fold Change Bacterial ncRNA
Absorbance (OD600nm)
CFU/ml
Time (mins)
Time (hrs)
-T2
+T2
T0
T20
T60
-log10 adjusted P-value
Log2 fold change
Translation and ribosomal regulation (J)
Transcription regulation (K)
DNA Replication, recombination and repair (L)
Cell cycle control and cell division (D)
Defense mechanisms (V)
Signal transduction mechanisms (T)
Cell wall/membrane/envelope (M)
Cell motility (N)
Extracellular structures (W)
Intracellular trafficking, secretion, transport (U)
Posttranslational modification and turnover (O)
Energy production and conversion (C)
Carbohydrate transport and metabolism (G)
Amino acid transport and metabolism (E)
Nucleotide transport and metabolism (F)
Coenzyme transport and metabolism (H)
Lipid transport and metabolism (I)
Inorganic ion transport and metabolism (P)
Secondary metabolite transport and metabolism (Q)
General function prediction only (R)
Function unknown (S)
No COG assignment (X)
Downregulated
Upregulated
Number of genes
KilS
Tff
SokX
SgrS
SibD
RnpB
RyfA
GlmY
SsrS
SokE
PspH
FnrS
SdsR
CsrB
DsrA
MotR
MicF
SraG

**Supplementary Figure 1.** (**A**) Graph showing growth of *E. coli* -/+ T2 phage (added at T0) from which samples for RIL/RNA-seq were taken (at T0, T20 and T60; indicated). (**B**) Graph showing the number of viable cells as a function of time in the T2 infected culture enumerated by colony forming units (CFU) linked to (A). (**C**) Graph showing the growth of *E. coli* -/+ T2 phage added at T0 over a 24 h period. Error bars represent standard error (n=3). (**D**) Volcano plot of differential RNA abundance at $T20^{INF}$ shown as a $\log_2$ fold-change from T20. Analysis performed by DESeq2. Differentially expressed genes (defined as those RNA with expression levels changed ≥ 2-fold with a false discovery rate adjusted p≤0.05) coloured as blue (bacterial) and orange (T2). Bacterial RNA whose abundance did not change significantly are shown in grey. (**E**) As (D) but comparing $T60^{INF}$ against T60. (**F**) Clustering of Orthologous Groups (COG) annotation of differentially expressed genes from (D) and (E). (**G**) As in (D) and (E) but showing differential non-coding RNA abundance at $T20^{INF}$ coloured in green and labelled. (**H**) As in (G) but comparing $T60^{INF}$ against T60.

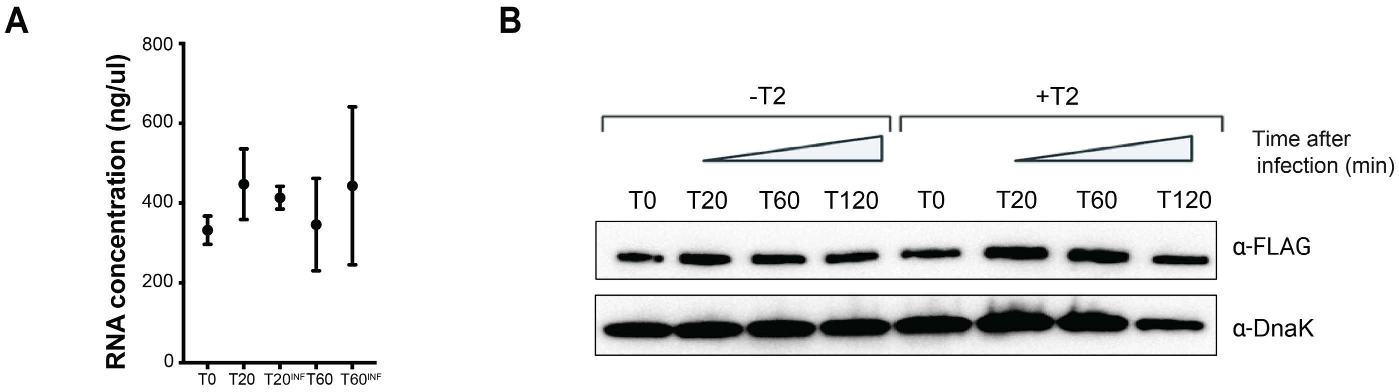


**Supplementary Figure 2.** (**A**) Graph showing the average concentration (n=3) of the total RNA sample collected for RIL/RNA-seq analysis at the indicated time points. Error bars represent standard error. (**B**) Representative immunoblot of whole cell extracts of *E. coli* -/+ at different time points (minutes) following T2 infection. T0 indicates the time point prior to T2 infection. The Immunoblots were probed with α-3XFLAG antibody to detect 3XFLAG-tagged Hfq and α-DnaK antibody to detect DnaK as the loading control.

| sRNA | mRNA | T0 | T60$^{INF}$ |
|---|---|---|---|
| ArcZ | *aspA* | 73 | 125 |
| | *flgG* | 41 | 39 |
| | *glpT* | 48 | 77 |
| | *gpsA* | 45 | 20 |
| | *hupB* | 32 | 13 |
| | *napF* | 2284 | 158 |
| | *ompA* | 27 | 35 |
| | *ompC* | 46 | 51 |
| | *ompF* | 25 | 48 |
| | *rbsD* | 20612 | 7574 |
| | *tar* | 50 | 187 |
| | *tnaA* | 57 | 202 |
| | *tnaC* | 134 | 5937 |
| | *ybdA_3* | 44 | 21 |
| | *ydfW* | 54 | 55 |
| | *ypfM* | 101 | 1028 |
| | *ytfR* | 59 | 43 |
| ChiX | *ansB* | 35 | 10 |
| | *bolA* | 109 | 13 |
| | *chbB* | 1387 | 66 |
| | *chiP* | 4733 | 511 |
| | *dmsA* | 31 | 13 |
| | *fliA* | 35 | 19 |
| | *fliC* | 338 | 113 |
| | *flxA* | 85 | 29 |
| | *gatZ* | 29 | 10 |
| | *glnA* | 43 | 15 |
| | *gltI* | 424 | 114 |
| | *gltJ* | 112 | 70 |
| | *gyrA* | 250 | 47 |
| | *hupB* | 1196 | 267 |
| | *infB* | 90 | 46 |
| | *miaA* | 31 | 14 |
| | *nlpD* | 279 | 36 |
| | *ompF* | 123 | 80 |
| | *raiA* | 46 | 23 |
| | *rbsD* | 83 | 26 |
| | *rplK* | 312 | 10 |
| | *rpoS* | 41 | 11 |
| | *rpsF* | 211 | 17 |
| | *sstT* | 343 | 52 |
| | *tar* | 31 | 12 |
| | *tdcA* | 33 | 11 |
| | *tdcG* | 148 | 11 |
| | *tmcA* | 25 | 12 |
| | *tnaA* | 26 | 16 |
| | *ytfK* | 96 | 24 |
| CpxQ | *dtpB* | 24 | 23 |
| | *glnA* | 54 | 23 |
| | *glpK* | 70 | 397 |
| | *glpT* | 39 | 98 |
| | *hupB* | 704 | 188 |
| | *hycA* | 24 | 47 |
| | *lon* | 156 | 71 |
| | *nrfA* | 59 | 12 |
| | *ompF* | 34 | 50 |
| | *pheT* | 41 | 65 |
| | *skp* | 71 | 32 |
| CyaR | *araJ* | 34 | 11 |
| | *cyoB* | 44 | 16 |
| | *dcuC* | 27 | 31 |
| | *gntU* | 557 | 49 |
| | *hfq* | 182 | 18 |
| | *kgtP* | 49 | 20 |
| | *miaA* | 167 | 16 |
| | *ompA* | 75 | 10 |
| | *ompX* | 1123 | 125 |
| | *ptsG* | 621 | 47 |
| | *rfbA* | 33 | 16 |
| | *rimP* | 53 | 10 |
| | *rpsT* | 157 | 13 |
| | *slyB* | 64 | 96 |
| | *tsx* | 51 | 14 |
| | *yebO* | 1641 | 768 |
| | *yecH* | 64 | 14 |
| | *yeiH* | 133 | 52 |
| | *yfdY* | 36 | 36 |
| DsrA | *gyrA* | 45 | 15 |
| | *nlpD* | 722 | 21 |
| FnrS | *ahpF* | 89 | 12 |
| | *dinI* | 79 | 59 |
| | *iscR* | 88 | 12 |
| | *marB* | 63 | 10 |
| | *ytfK* | 274 | 604 |
| GcvB | *argT* | 1138 | 493 |
| | *dinD* | 43 | 12 |
| | *dppA* | 143 | 10 |
| | *fliC* | 356 | 35 |
| | *flxA* | 161 | 10 |
| | *gatY* | 36 | 14 |
| | *gdhA* | 807 | 68 |
| | *gltI* | 3294 | 185 |
| | *gltJ* | 5286 | 910 |
| | *gyrA* | 96 | 10 |
| | *ompF* | 147 | 17 |
| | *raiA* | 248 | 50 |
| | *sstT* | 3477 | 1207 |
| | *tcyJ* | 155 | 22 |
| | *ydhC* | 66 | 38 |
| | *yifK* | 940 | 12 |
| GlmZ | *glmS* | 10363 | 1744 |
| MicA | *hupB* | 23 | 493 |
| | *ompA* | 432 | 12 |
| | *tolB* | 156 | 10 |

| sRNA | mRNA | T0 | T60$^{INF}$ |
|---|---|---|---|
| MicL | *acpP* | 169 | 41 |
| | *lpp* | 66 | 83 |
| MicF | *ompF* | 637 | 1047 |
| MgrR | *eptB* | 885 | 86 |
| | *fliC* | 1100 | 87 |
| | *gcvT* | 139 | 11 |
| | *htrL* | 79 | 24 |
| | *insA-7* | 343 | 43 |
| | *yebZ* | 141 | 28 |
| | *ygdQ* | 465 | 14 |
| | *ygfZ* | 63 | 18 |
| OxyS | *glpF* | 145 | 126 |
| RprA | *ompF* | 54 | 35 |
| | *ymgC* | 104 | 60 |
| RybB | *dmsA* | 428 | 29 |
| | *rbsK* | 28 | 14 |
| RydC | *gyrA* | 34 | 26 |

| mRNA | mRNA | T0 | T60$^{INF}$ |
|---|---|---|---|
| *cpsA* | *ydfH* | 40 | 140 |
| *fdoG* | *sucD* | 45 | 40 |
| *flgL* | *chiP* | 18 | 13 |
| | *cof* | 24 | 80 |
| | *fliC* | 36 | 86 |
| | *gltI* | 31 | 220 |
| | *gltJ* | 28 | 342 |
| | *gyrA* | 24 | 189 |
| | *hupB* | 25 | 44 |
| | *ompA* | 14 | 14 |
| | *rplL* | 12 | 10 |
| | *sucD* | 13 | 76 |
| | *ytfK* | 20 | 18 |
| *fliC* | *flhC* | 30 | 26 |
| | *fliA* | 541 | 293 |
| | *gltJ* | 17 | 12 |
| | *uhpT* | 40 | 13 |
| *fliA* | *mglB* | 10 | 60 |
| | *raiA* | 27 | 33 |
| | *rpsP* | 24 | 150 |
| | *rpsS* | 10 | 66 |
| *glnA* | *sucA* | 32 | 16 |
| *glpK* | *glpX* | 33 | 11 |
| *gltJ* | *gyrA* | 10 | 47 |
| *gyrA* | *uhpT* | 10 | 55 |
| *hemG* | *murI* | 178 | 14 |
| *insA-2* | *insA-5* | 108 | 11 |
| *insA-2* | *insA-6* | 82 | 31 |
| *insA9* | *insA-5* | 78 | 32 |
| | *insA-6* | 195 | 41 |
| *insB-2* | *insB-6* | 37 | 32 |
| *insB-3* | *insB-5* | 41 | 28 |
| *insB-4* | *insB9* | 53 | 22 |
| *insH-10* | *insH-8* | 263 | 95 |
| *insH-11* | *insH-7* | 310 | 61 |
| *insH-2* | *insH21* | 422 | 88 |
| *insH-3* | *insH-4* | 414 | 87 |
| *insH-9* | *insH-5* | 78 | 16 |
| | *insH-6* | 282 | 63 |
| *insI-1* | *insI-2* | 64 | 10 |
| *insL-1* | *insL-2* | 222 | 15 |
| *minD* | *ykgH* | 18 | 20 |
| *ompC* | *uhpT* | 14 | 10 |
| *ompF* | *mdtK* | 12 | 13 |
| *ompF* | *ybiJ* | 20 | 32 |
| *rpsG* | *uhpT* | 17 | 52 |
| *tufA* | *tufB* | 2726 | 837 |
| *ykgH* | *wzzB* | 30 | 13 |
| | *ynaM* | 23 | 18 |

| sRNA | sRNA | T0 | T60$^{INF}$ |
|---|---|---|---|
| ArcZ | ChiX | 131 | 40 |
| | CpxQ | 10 | 26 |
| | CyaR | 20 | 21 |
| | FlgO | 33 | 115 |
| | FnrS | 35 | 54 |
| | GadY | 14 | 20 |
| | GcvB | 27 | 13 |
| | MgrR | 10 | 12 |
| | OmrA | 18 | 26 |
| | OmrB | 24 | 19 |
| | Spf | 31 | 10 |
| ChiX | CpxQ | 86 | 25 |
| | CyaR | 103 | 33 |
| | DsrA | 614 | 39 |
| | FlgO | 404 | 3570 |
| | FnrS | 349 | 57 |
| | GadY | 78 | 14 |
| | GcvB | 1378 | 136 |
| | GlmZ | 678 | 122 |
| | McaS | 105 | 121 |
| | MgrR | 490 | 17 |
| | MicL | 202 | 15 |
| | OmrA | 61 | 37 |
| | OxyS | 147 | 58 |
| | RybB | 29 | 22 |
| | RydC | 49 | 20 |
| | Spf | 1498 | 14 |
| | SucD | 55 | 163 |
| | UhpU | 73 | 11 |
| CyaR | FlgO | 38 | 33 |
| | GcvB | 61 | 12 |
| FlgO | GcvB | 92 | 57 |
| | MgrR | 29 | 13 |
| GcvB | MgrR | 133 | 22 |

**Supplementary Figure 3.** Table showing a subset of chimera counts from RIL-seq analysis of RNA-RNA interactions between bacterial mRNA-mRNA, sRNA-mRNA and sRNA-sRNA which were present at T0 and $T60^{INF}$ time points and therefore deemed 'core' interactions.

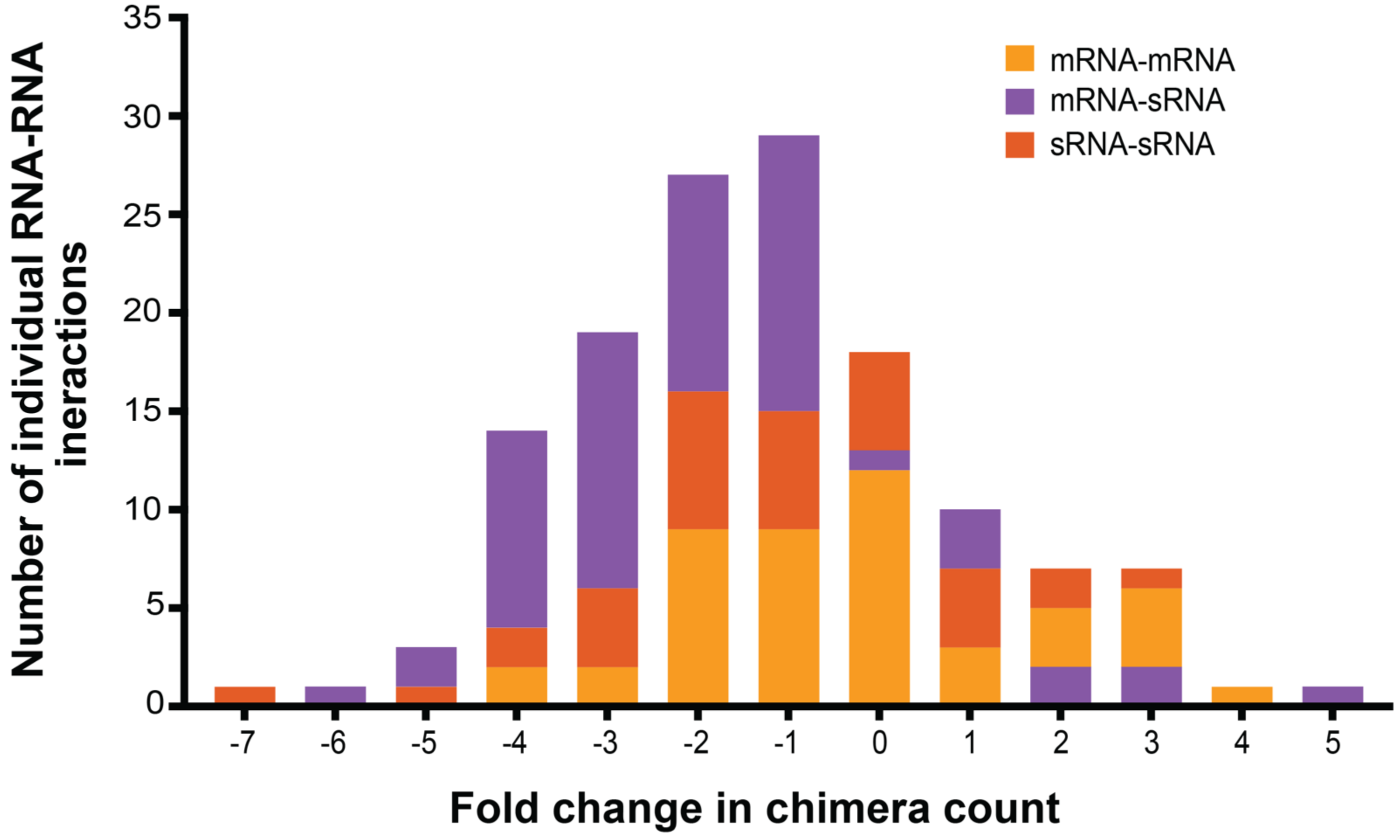


**Supplementary Figure 4.** Overlayed bar chart showing the total number of each chimera type (mRNA-mRNA, mRNA-sRNA, sRNA-sRNA) which constitute the core interactions and their relative fold-change between time point T0 and $T60^{INF}$. The different types of chimeric interactions are grouped, and colour-coded as follows: mRNA-mRNA (yellow), mRNA-sRNA (purple), sRNA-sRNA (orange).

| sRNA | mRNA | T0 | $T60^{INF}$ |
|---|---|---|---|
| ArcZ | *aspA* | 73 | 125 |
| | *flgG* | 41 | 39 |
| | *glpT* | 48 | 77 |
| | *gpsA* | 45 | 20 |
| | *hupB* | 32 | 13 |
| | *napF* | 2284 | 158 |
| | *ompA* | 27 | 35 |
| | *ompC* | 46 | 51 |
| | *ompF* | 25 | 48 |
| | *rbsD* | 20612 | 7574 |
| | *tar* | 50 | 187 |
| | *tnaA* | 57 | 202 |
| | *tnaC* | 134 | 5937 |
| | *ybdA_3* | 44 | 21 |
| | *ydfW* | 54 | 55 |
| | *ypfM* | 101 | 1028 |
| | *ytfR* | 59 | 43 |
| ChiX | *ansB* | 35 | 10 |
| | *bolA* | 109 | 13 |
| | *chbB* | 1387 | 66 |
| | *chiP* | 4733 | 511 |
| | *dmsA* | 31 | 13 |
| | *fliA* | 35 | 19 |
| | *fliC* | 338 | 113 |
| | *flxA* | 85 | 29 |
| | *gatZ* | 29 | 10 |
| | *glnA* | 43 | 15 |
| | *gltI* | 424 | 114 |
| | *gltJ* | 112 | 70 |
| | *gyrA* | 250 | 47 |
| | *hupB* | 1196 | 267 |
| | *infB* | 90 | 46 |
| | *miaA* | 31 | 14 |
| | *nlpD* | 279 | 36 |
| | *ompF* | 123 | 80 |
| | *raiA* | 46 | 23 |
| | *rbsD* | 83 | 26 |
| | *rplK* | 312 | 10 |
| | *rpoS* | 41 | 11 |
| | *rpsF* | 211 | 17 |
| | *sstT* | 343 | 52 |
| | *tar* | 31 | 12 |
| | *tdcA* | 33 | 11 |
| | *tdcG* | 148 | 11 |
| | *tmcA* | 25 | 12 |
| | *tnaA* | 26 | 16 |
| | *ytfK* | 96 | 24 |
| CpxQ | *dtpB* | 24 | 23 |
| | *glnA* | 54 | 23 |
| | *glpK* | 70 | 397 |
| | *glpT* | 39 | 98 |
| | *hupB* | 704 | 188 |
| | *hycA* | 24 | 47 |
| | *lon* | 156 | 71 |
| | *nrfA* | 59 | 12 |
| | *ompF* | 34 | 50 |
| | *pheT* | 41 | 65 |
| | *skp* | 71 | 32 |
| CyaR | *araJ* | 34 | 11 |
| | *cyoB* | 44 | 16 |
| | *dcuC* | 27 | 31 |
| | *gntU* | 557 | 49 |
| | *hfq* | 182 | 18 |
| | *kgtP* | 49 | 20 |
| | *miaA* | 167 | 16 |
| | *ompA* | 75 | 10 |
| | *ompX* | 1123 | 125 |
| | *ptsG* | 621 | 47 |
| | *rfbA* | 33 | 16 |
| | *rimP* | 53 | 10 |
| | *rpsT* | 157 | 13 |
| | *slyB* | 64 | 96 |
| | *tsx* | 51 | 14 |
| | *yebO* | 1641 | 768 |
| | *yecH* | 64 | 14 |
| | *yeiH* | 133 | 52 |
| | *yfdY* | 36 | 36 |
| DsrA | *gyrA* | 45 | 15 |
| | *nlpD* | 722 | 21 |
| FnrS | *ahpF* | 89 | 12 |
| | *dinI* | 79 | 59 |
| | *iscR* | 88 | 12 |
| | *marB* | 63 | 10 |
| | *ytfK* | 274 | 604 |
| GcvB | *argT* | 1138 | 493 |
| | *dinD* | 43 | 12 |
| | *dppA* | 143 | 10 |
| | *fliC* | 356 | 35 |
| | *flxA* | 161 | 10 |
| | *gatY* | 36 | 14 |
| | *gdhA* | 807 | 68 |
| | *gltI* | 3294 | 185 |
| | *gltJ* | 5286 | 910 |
| | *gyrA* | 96 | 10 |
| | *ompF* | 147 | 17 |
| | *raiA* | 248 | 50 |
| | *sstT* | 3477 | 1207 |
| | *tcyJ* | 155 | 22 |
| | *ydhC* | 66 | 38 |
| | *yifK* | 940 | 12 |
| GlmZ | *glmS* | 10363 | 1744 |
| MicA | *hupB* | 23 | 493 |
| | *ompA* | 432 | 12 |
| | *tolB* | 156 | 10 |

| sRNA | mRNA | T0 | $T60^{INF}$ |
|---|---|---|---|
| MicL | *acpP* | 169 | 41 |
| | *lpp* | 66 | 83 |
| MicF | *ompF* | 637 | 1047 |
| MgrR | *eptB* | 885 | 86 |
| | *fliC* | 1100 | 87 |
| | *gcvT* | 139 | 11 |
| | *htrL* | 79 | 24 |
| | *insA-7* | 343 | 43 |
| | *yebZ* | 141 | 28 |
| | *ygdQ* | 465 | 14 |
| | *ygfZ* | 63 | 18 |
| OxyS | *glpF* | 145 | 126 |
| RprA | *ompF* | 54 | 35 |
| | *ymgC* | 104 | 60 |
| RybB | *dmsA* | 428 | 29 |
| | *rbsK* | 28 | 14 |
| RydC | *gyrA* | 34 | 26 |

| mRNA | mRNA | T0 | $T60^{INF}$ |
|---|---|---|---|
| *cpsA* | *ydfH* | 40 | 140 |
| *fdoG* | *sucD* | 45 | 40 |
| *flgL* | *chiP* | 18 | 13 |
| | *cof* | 24 | 80 |
| | *fliC* | 36 | 86 |
| | *gltI* | 31 | 220 |
| | *gltJ* | 28 | 342 |
| | *gyrA* | 24 | 189 |
| | *hupB* | 25 | 44 |
| | *ompA* | 14 | 14 |
| | *rplL* | 12 | 10 |
| | *sucD* | 13 | 76 |
| | *ytfK* | 20 | 18 |
| *fliC* | *flhC* | 30 | 26 |
| | *fliA* | 541 | 293 |
| | *gltJ* | 17 | 12 |
| | *uhpT* | 40 | 13 |
| *fliA* | *mglB* | 10 | 60 |
| | *raiA* | 27 | 33 |
| | *rpsP* | 24 | 150 |
| | *rpsS* | 10 | 66 |
| *glnA* | *sucA* | 32 | 16 |
| *glpK* | *glpX* | 33 | 11 |
| *gltJ* | *gyrA* | 10 | 47 |
| *gyrA* | *uhpT* | 10 | 55 |
| *hemG* | *murI* | 178 | 14 |
| *insA-2* | *insA-5* | 108 | 11 |
| *insA-2* | *insA-6* | 82 | 31 |
| *insA9* | *insA-5* | 78 | 32 |
| | *insA-6* | 195 | 41 |
| *insB-2* | *insB-6* | 37 | 32 |
| *insB-3* | *insB-5* | 41 | 28 |
| *insB-4* | *insB9* | 53 | 22 |
| *insH-10* | *insH-8* | 263 | 95 |
| *insH-11* | *insH-7* | 310 | 61 |
| *insH-2* | *insH21* | 422 | 88 |
| *insH-3* | *insH-4* | 414 | 87 |
| *insH-9* | *insH-5* | 78 | 16 |
| | *insH-6* | 282 | 63 |
| *insI-1* | *insI-2* | 64 | 10 |
| *insL-1* | *insL-2* | 222 | 15 |
| *minD* | *ykgH* | 18 | 20 |
| *ompC* | *uhpT* | 14 | 10 |
| *ompF* | *mdtK* | 12 | 13 |
| *ompF* | *ybiJ* | 20 | 32 |
| *rpsG* | *uhpT* | 17 | 52 |
| *tufA* | *tufB* | 2726 | 837 |
| *ykgH* | *wzzB* | 30 | 13 |
| | *ynaM* | 23 | 18 |

| sRNA | sRNA | T0 | $T60^{INF}$ |
|---|---|---|---|
| ArcZ | ChiX | 131 | 40 |
| | CpxQ | 10 | 26 |
| | CyaR | 20 | 21 |
| | FlgO | 33 | 115 |
| | FnrS | 35 | 54 |
| | GadY | 14 | 20 |
| | GcvB | 27 | 13 |
| | MgrR | 10 | 12 |
| | OmrA | 18 | 26 |
| | OmrB | 24 | 19 |
| | Spf | 31 | 10 |
| ChiX | CpxQ | 86 | 25 |
| | CyaR | 103 | 33 |
| | DsrA | 614 | 39 |
| | FlgO | 404 | 3570 |
| | FnrS | 349 | 57 |
| | GadY | 78 | 14 |
| | GcvB | 1378 | 136 |
| | GlmZ | 678 | 122 |
| | McaS | 105 | 121 |
| | MgrR | 490 | 17 |
| | MicL | 202 | 15 |
| | OmrA | 61 | 37 |
| | OxyS | 147 | 58 |
| | RybB | 29 | 22 |
| | RydC | 49 | 20 |
| | Spf | 1498 | 14 |
| | SucD | 55 | 163 |
| | UhpU | 73 | 11 |
| CyaR | FlgO | 38 | 33 |
| | GcvB | 61 | 12 |
| FlgO | GcvB | 92 | 57 |
| | MgrR | 29 | 13 |
| GcvB | MgrR | 133 | 22 |

**Supplementary Figure 5.** Table showing chimeras between T2-RNA and a bacterial sRNA or mRNA. Presence of each RNA-RNA interaction at the two infection time points ($T20^{INF}$ and $T60^{INF}$) is indicated.

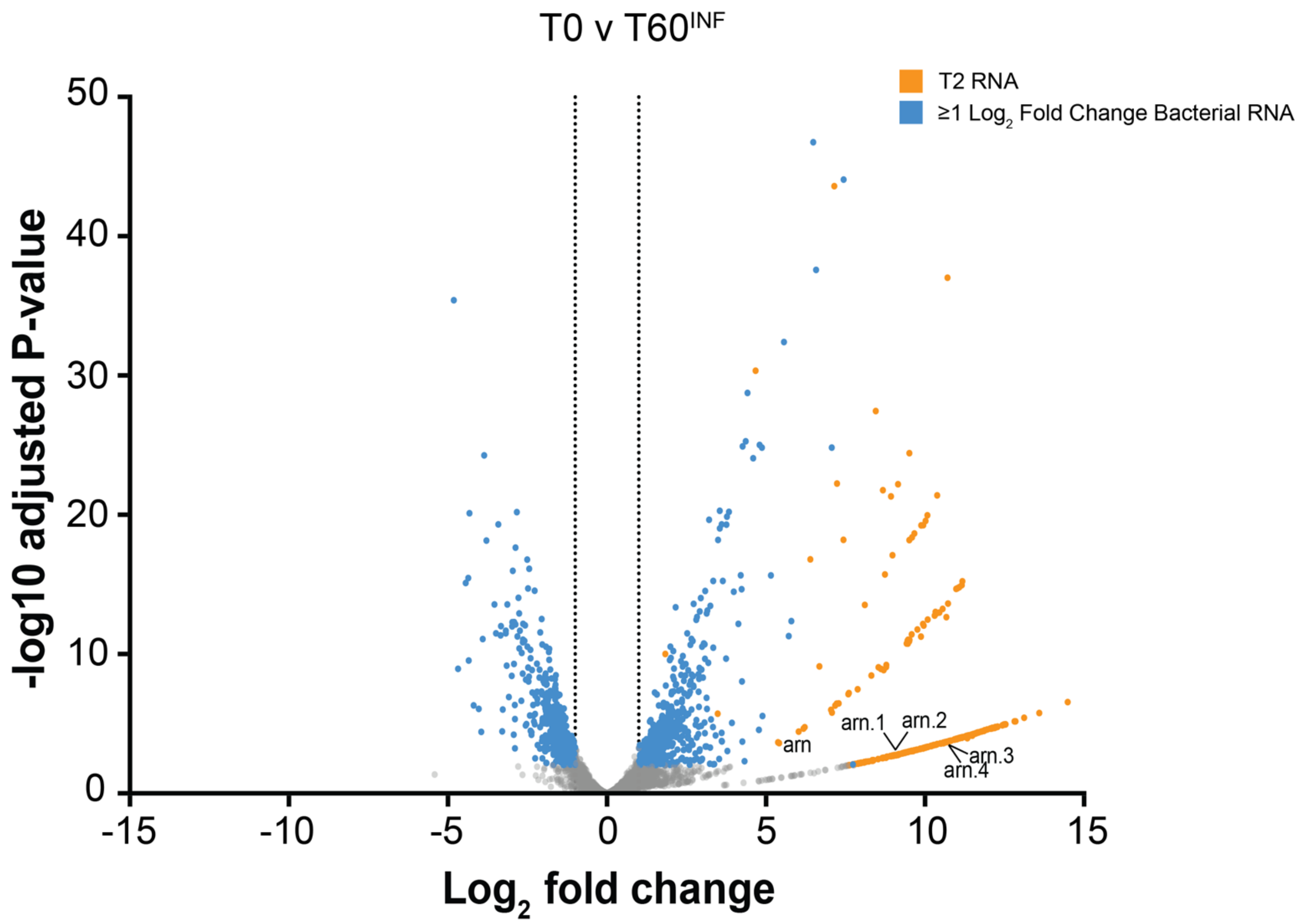


**Supplementary Figure 6.** Volcano plot of differential RNA abundance at $T60^{INF}$ shown as a $log_2$ fold-change from T0. Analysis performed by DESeq2. Differentially expressed genes (defined as those RNA with expression levels changed ≥ 2-fold with a adjusted false discovery rate p≤0.05) coloured as blue (bacterial) and orange (T2). Bacterial RNA whose abundance did not change significantly are shown in grey. The *arn* genes are indicated.

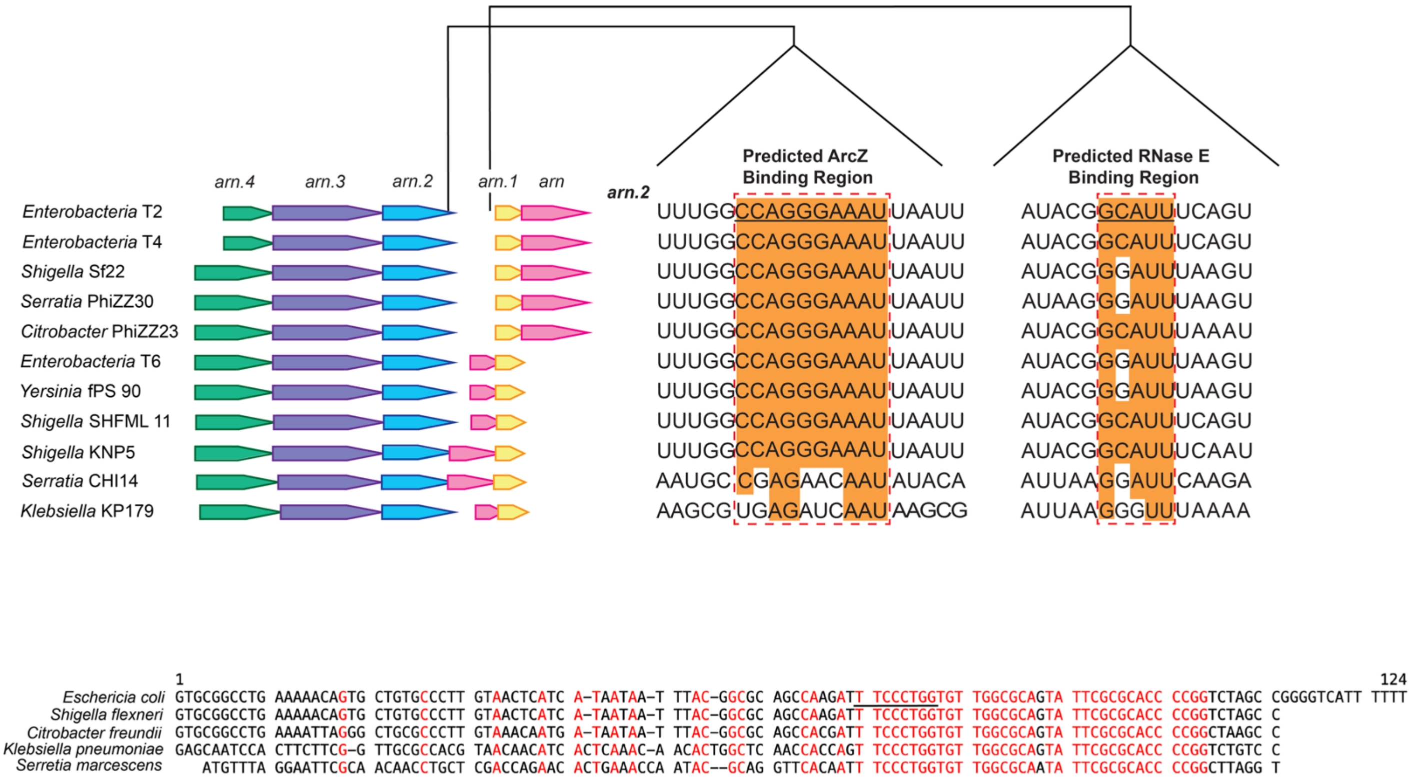


**Supplementary Figure 7.** Schematic representing the *arn* operon in different T2-like phages obtained using WebFlaGs (50). The predicted ArcZ and RNase E binding regions are aligned and conserved residues highlighted (in orange). Sequence alignment of ArcZ from each host is shown at the bottom, with conserved residues highlighted in red, and the region that interacts with the T2 *arn* transcript underlined.

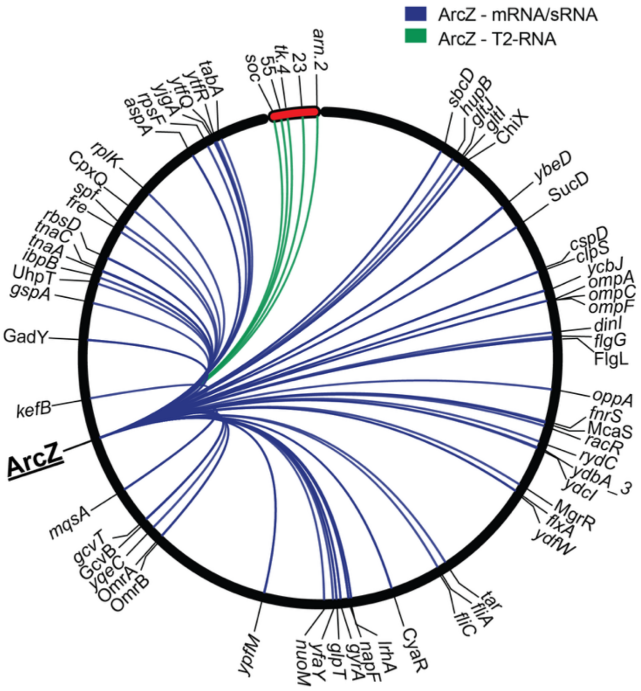


| | T0 | T20 | T20[INF] | T60 | T60[INF] |
|---|---|---|---|---|---|
| ArcZ-mRNA | 65 | 282 | 315 | 291 | 49 |
| ArcZ-sRNA | 12 | 21 | 22 | 23 | 14 |
| ArcZ-T2-RNA | 0 | 0 | 5 | 0 | 4 |

**Supplementary Figure 8.** Circos plot of ArcZ-RNA interactions, represented by at least 10 chimeric fragments, mapped to the *E. coli* MG1655 and T2 phage genomes. The T2 genome is indicated as a red bar at the 12 o'clock position. The different types of ArcZ-RNA interactions are coloured blue (ArcZ interactions with bacterial RNA) and green (ArcZ interactions with T2 RNA). Number of unique interactions between ArcZ and either a bacterial mRNA or sRNA, or T2-RNA at each time point are indicated.
