## Supplementary Data 1 for "A nucleic acid host factor enables optimal phage replication in *Escherichia coli*"

| <u>Name</u> | <u>Description</u> | <u>Source or Reference</u> |
|---|---|---|
| WT | *E. coli* MG1655 | *E. coli* Genetic Stock Centre |
| *hfq*::3XFLAG | *E. coli* MG1655 *hfq-3XFLAG-kan* | Gift by Prof. Jörg Vogel, also (McQuail *et al.* 2024) |
| Δ*hfq* | *E. coli* BW25113 Δ*hfq-kan* | *E. coli* Genetic Stock Centre |
| Δ*arcZ* | *E. coli* MG1655 Δ*arcZ-zeo* | Gift by Dr Susan Gottesman |
| Δ*rpoS* | *E. coli* BW25113 Δ*rpoS-kan* | (Baba *et al.* 2006) |
| Δ*mcrC* | *E. coli* BW25113 Δ*mcrC-kan* | (Baba *et al.* 2006) |
| Δ*arcZ*Δ*mcrC* | *E. coli* BW25113 Δ*arcZ*Δ*mcrC-kan* | This study |
| **Plasmids** | | |
| pBAD24 | Plasmid containing an L-arabinose inducible promoter (*$bla_{TEM-1}$*) | (Guzman *et al.* 1995) |
| pHfq | pBAD24::*hfq*-FLAG | Gift by Prof. Jörg Vogel; also (McQuail *et al.* 2025) |
| pHfq Q8A | pBAD24::*hfq*-FLAG with Q8A mutation | (McQuail et al. 2025) |
| pHfq D9A | pBAD24::*hfq* with D9A mutation | (McQuail et al. 2025) |
| pHfq R16A | pBAD24::*hfq* with R16A mutation | (McQuail et al. 2025) |
| pHfq R17A | pBAD24::*hfq* with R17A mutation | (McQuail et al. 2025) |
| pHfq Y25A | pBAD24::*hfq* with Y25A mutation | (McQuail et al. 2025) |
| pHfq ΔCTD | pBAD24::*hfq* with C-terminal binding domain (CTD) deletion | (McQuail et al. 2025) |
| pKF68-3 | ColE1 plasmid based on pZE12-luc; expresses Salmonella SdsR from IPTG-inducible $PL_{lacO}$ promoter (*$bla_{TEM-1}$*) | Gift by Prof. Jörg Vogel; also (Frohlich *et al.* 2016) |
| pArcZ | pKF68-3 with *Salmonella* SdsR changed to *E. coli* ArcZ (*$bla_{TEM-1}$*) | This study |
| pBAD18 | Plasmid containing an L-arabinose inducible promoter(*$bla_{TEM-1}$*) | ThermoFisher |
| Construct I | pBAD18 with the nucleotide sequence corresponding *arn.2* and IR region with a 5' sequence encoding a 6His-tag under pBAD promoter | This study |
| Construct II | pBAD18 with the nucleotide sequence corresponding *arn.2* with a 5' sequence encoding a 6His-tag under pBAD promoter | This study |
| Construct III | pBAD18 with the nucleotide sequence corresponding *arn.2* and truncated IR region with a 5' sequence encoding a 6His-tag under pBAD promoter | This study |

| | | |
|---|---|---|
| Construct IV | pBAD18 with the nucleotide sequence corresponding the entire *arn* operon with a 3’ sequence encoding a 6His-tag under pBAD promoter | This study |
| Construct V | pBAD18 with the nucleotide sequence corresponding the entire *arn* operon with the RNase E binding region in the IR mutated and with a 3’ sequence encoding a 6His-tag under pBAD promoter | This study |
